# The circadian clock conveys thermal and photoperiodic cues to modulate EYES ABSENT via the neuropeptide PDF to regulate seasonal physiology

**DOI:** 10.1101/2022.10.27.514061

**Authors:** Sergio Hidalgo, Maribel Anguiano, Christine A. Tabuloc, Joanna C. Chiu

## Abstract

Organisms adapt to seasonal changes in photoperiod and temperature to survive; however, the mechanisms by which these signals are integrated in the brain are poorly understood. We previously reported that EYES ABSENT (EYA) in *Drosophila* shows higher levels in cold temperature or short photoperiod, and genetic ablation of *eya* in the fly brain inhibits reproductive dormancy, suggesting that EYA promotes winter physiology. Nevertheless, the mechanisms by which EYA senses seasonal cues are unclear. Pigment-Dispersing Factor (PDF) is a neuropeptide important for photoentrainment and regulation of circadian output rhythms. Interestingly, PDF also regulates reproductive dormancy, suggesting that it may mediate the function of the circadian clock in modulating seasonal physiology. In this study, we investigated the role of PDF signaling in mediating the impact of EYA on seasonal biology. First, we subjected flies to different photoperiodic and temperature regimes and observed that PDF abundance is lower in cold and short days, compared to warm and long days. Interestingly, the response of PDF to seasonal cues is opposite of what was observed for EYA. We then determined the potential for PDF to convey seasonal cues and modulate EYA function in seasonality by assessing coexpression of EYA and PDF receptor. Our results indicated that PDF receptor (PDFR) is indeed coexpressed with EYA in the fly brain, including in the circadian clock neuronal network and neurons in the *pars intercerebralis*. We then manipulated PDF signaling in *eya*+ cells to show that PDF modulates seasonal adaptations in daily activity rhythm and ovary development via EYA-dependent and independent mechanisms. At the molecular level, manipulating PDF signaling impacted EYA protein abundance. Specifically, we showed that protein kinase A (PKA), an effector of PDF signaling, phosphorylates EYA and promotes its degradation. This explains the opposite responses of PDF and EYA abundance to changes in seasonal cues. In summary, our results support a model in which PDF signaling negatively modulates EYA levels to regulate seasonal physiology, linking the circadian clock to the modulation of seasonal adaptations.

## Introduction

As seasons change, organisms adapt to adverse environmental conditions to survive^1,2^. While some animals, like monarch butterflies^3,4^, migrate into more favorable environments, others undergo metabolic and physiological changes that allow them to prevail through the harsh winter^5,6^. These seasonal adaptations have been described in plants^6,7^ and animals^8–11^, and are triggered by changes in photoperiod^12,13^ (i.e. daylength) and temperature^14^. However, the molecular mechanisms that allow organisms to extract environmental information reflecting the calendar year is not well established. This is especially true in animals.

In several animals, endocrine signals drive seasonal adaptations. In mammals and birds, the increases in the hormone thyrotrophin (TSH) and the downstream T3 hormone, are key for photoperiodic summer responses^15–18^. In insects, reduction of insulin-like peptides (ILPs) and juvenile hormone (JH) are important for overwintering^19–22^. Although the hormonal components differ between species, studies have suggested that EYES ABSENT (EYA), a cotranscription factor with phosphatase activity, could be a key conserved molecular player within the seasonal timer, acting upstream of endocrine responses to modulate seasonal physiology. In the Japanese quail (*Coturnix japonica*), the expression of the *eya* homolog, *Eya3*, increases in the *pars tuberalis* of the pituitary gland upon transitioning into summer^17^. This phenomenon is also observed in soay sheep (*Ovis aries*), where EYA3 protein has also been proposed as an effector upstream TSH-β^18,23^. We recently described a functional role of EYA in the seasonal response of *Drosophila melanogaster*^24^. Specifically, we showed that *eya* expression in the insulin-producing cells (IPCs) within the *pars intercerebralis* (*PI*), the insect ortholog of the hypothalamus, is critical for reproductive dormancy, a seasonal adaptation characterized by a reduction in ovary size and egg production. Moreover, we reported that *eya* mRNA and EYA protein levels are sensitive to changes in photoperiod and temperature, increasing in simulated winter conditions^24^. Despite evidence from multiple organisms, the mechanism mediating changes in EYA expression in response to photoperiod and temperature has yet to be determined.

The Bünning hypothesis states that the circadian clock is required for appropriate photoperiodic measurement^25,26^. The core clock in *Drosophila* is a transcriptional translational feedback loop (TTFL) composed of the positive elements CLK-CYC, which promote the transcription of the negative elements *period* (*per*) and *timeless* (*tim*) as well as other clock-regulated genes^27^. Accumulation and subsequent nuclear translocation of PER-TIM then leads to the repression of CLK-CYC activity, which in turn shuts down the transcription of *per* and *tim*, preparing the clock for the next cycle. The proteasome mediates light-dependent and light-independent degradation of the negative elements, ensuring the proper functioning and entrainment of the TTFL. Bünning proposed that circadian oscillations are composed of light and dark-requiring phases^26^. In short days, light is only present in the light phase, while in long days, light is now present in both light and in part of the dark-requiring phase. This was later modified and presented as “coincidence model”^28^. This model then assumes that the elements of the circadian clock are required for photoperiodism and seasonal adaptations. Studies in several insects have been instrumental in establishing this relationship^29–32^. Recently, Wood et al. proposed a mechanism in mammals where *Eya3* transcription is modulated by BMAL2-dependent chromatin remodeling in the *pars tuberalis* of the pituitary gland in sheep, highlighting the role of a circadian clock protein in regulating *Eya3* expression^33^. In *Drosophila* however, the IPCs do not express a functional circadian clock^34,35^, although TIM expression in the fly brain is found to modulate EYA abundance^24^. This suggests that other unknown clock mechanisms must be involved in relaying environmental information to EYA-expressing cells to modulate EYA function in seasonal adaptations.

The circadian clock neuronal network (CCNN) provides extensive connections to and modulates the IPCs^34–36^. The CCNN is composed of around 7 neuronal clusters: dorsal neurons-1 (DN1), dorsal neurons-2 (DN2), dorsal neurons-3 (DN3), lateral posterior neurons (LPN), dorsal lateral neurons (LNd), and the large and small ventral lateral neurons (l-LNvs and s-LNvs, respectively). Among CCNN clusters, DN1 and s-LNvs are known to modulate the IPCs^37–39^. The s-LNvs secretes Pigment Dispersing Factor (PDF), a neuropeptide that enables the molecular synchronization of neuronal clusters within the CCNN^40,41^, and is critical for the maintenance of anticipatory activity in light-dark cycles (LD) and the circadian rhythmicity in constant darkness (DD)^42–45^. Interestingly, the role of PDF in modulating seasonal adaptations has been examined in several organisms. In the blow fly and in the bean bug, ablation of the brain region containing the PDF-producing neurons prevents the discrimination between short and long photoperiod^46,47^. Moreover, ablation of *pdf* in the brown-winged green bug impairs photoperiodic control of reproduction^48^. In *Drosophila melanogaster*, photoperiod sensitivity is lost in *pdf* mutants^49^, whereas artificially increasing or decreasing the activity of *pdf*-expressing cells inhibits or promotes reproductive dormancy, respectively^50^. More importantly, the role of PDF in reproductive dormancy appears to be mediated by direct modulation of the IPCs^50^.

Here, we explore the relationship between PDF and EYA in the context of seasonal biology. We provide evidence to support a mechanism by which PDF signaling regulates multiple aspects of seasonal physiology by conveying environmental cues to alter EYA levels in the fly brain via a PKA-dependent post-translational mechanism. We proposed that during summer-like conditions, PDF promotes EYA degradation by PKA-mediated phosphorylation, while a reduction in *pdf* mRNA and PDF peptide in winter allows EYA accumulation, promoting winter adaptations.

## Results

### PDF levels change with photoperiod and temperature

The circadian neuropeptide PDF is known to exhibit daily cycling in abundance in the fly brain and can be synchronized by light^51^. Furthermore, *pdf* mRNA levels are sensitive to light intensity^52^. This suggests that PDF could be sensitive to environmental cues that change with seasons, namely photoperiod and temperature, and can subsequently convey these seasonal cues to alter EYA function in the fly brain. To test this hypothesis, we first examined the sensitivity of PDF abundance to key seasonal cues. We measured PDF levels in the dorsal terminals of the s-LNvs in flies subjected to different photoperiods and temperature conditions. First, we tested flies that were subjected to 7 days in simulated summer conditions, i.e. long photoperiod (LP, 16:8 LD) at 25°C, or simulated winter, i.e. short photoperiod (SP, 8:16 LD) at 10°C (Figure 1A-B). PDF levels showed daily rhythmicity in both conditions with peak phase at ZT0 (Rhythmicity LP25 p =5.4×10^−3^, Rhythmicity SP10 p = 3.19×10^−7^). Although we did not observe a difference in the amplitude of the PDF cycling (p = 0.22), a reduction in the Midline Estimating Statistic Of Rhythm (MESOR) in simulated winter compared to simulated summer was observed, suggesting an overall reduction of PDF levels in winter-like conditions (MESOR LP25 = 2.41, MESOR SP10 = 1.26, p = 2.06×10^−17^; Figure 1C).

**Figure 1:**
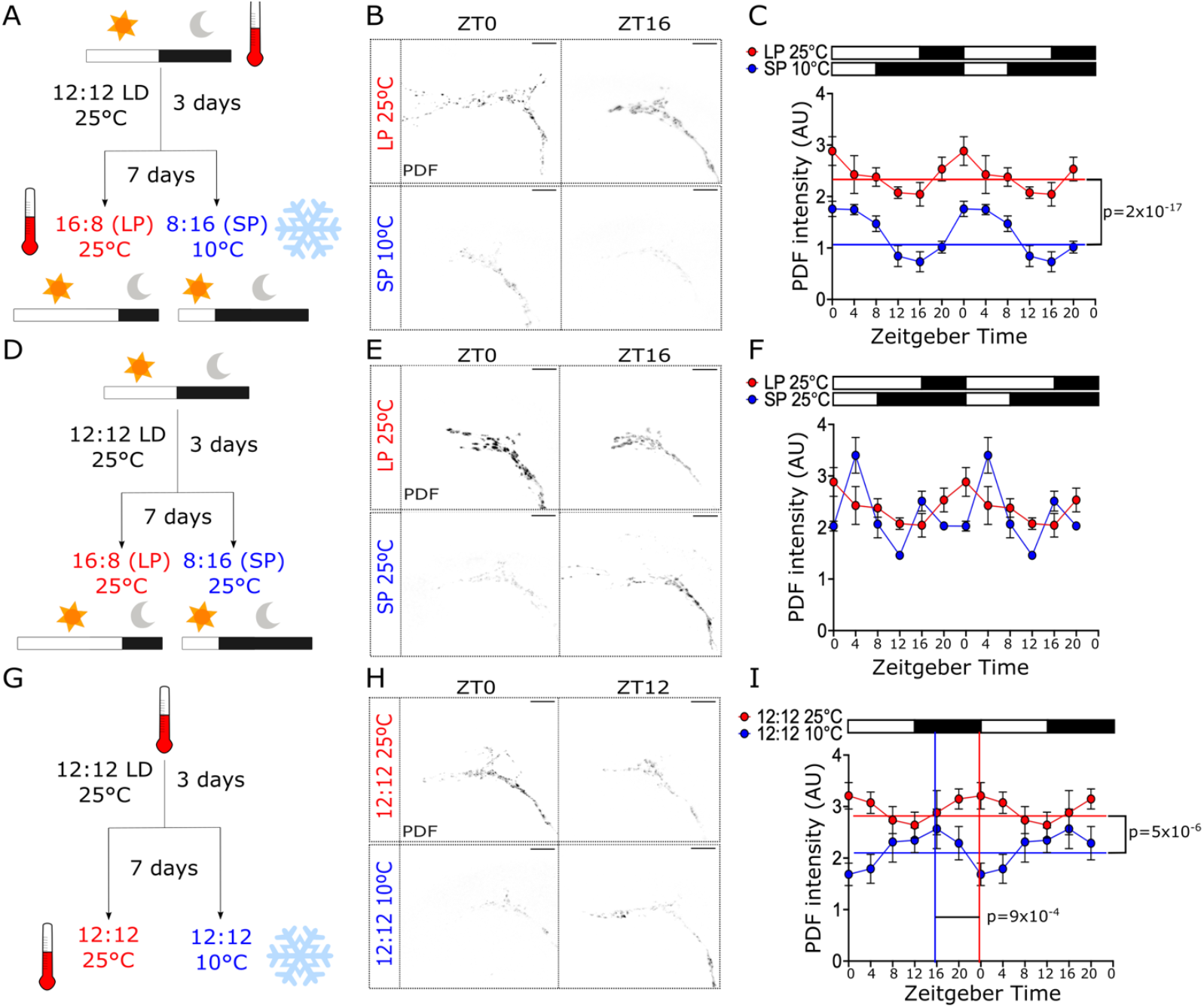
PDF levels are regulated by photoperiod and temperature. PDF levels in the s-LNv dorsal terminals were analyzed in flies entrained to (A) long-photoperiod (LP, 16:8 LD) at 25ºC or short-photoperiod (SP, 8:16 LD) at 10ºC, (D) LP or SP at 25ºC, and (G) 12:12 LD cycles at 25ºC or 10ºC. Representative images for these conditions at ZT0 and ZT16 or ZT12 are shown in B, E and H, respectively. Double-plotted graphs of PDF signal for each condition are shown in C, F and I, respectively. White boxes represent lights-on, and black boxes represent lights-off. Circacompare was used to determine rhythmicity. Differences in MESOR (horizontal lines) and phase (vertical lines) are displayed in the graphs if statistically significant (p < 0.05). Scale bars are 10 μm. Number of brains imaged were LP25: ZT0 n = 18, ZT4 n = 11, ZT8 n = 13, ZT12 n = 18, ZT16 = 9, ZT20 n = 14; SP10: ZT0 n = 9, ZT4 n = 8, ZT8 n = 14, ZT12 n = 14, ZT16 = 18, ZT20 n = 20; SP25 ZT0 n = 6, ZT4 n = 8, ZT8 n = 9, ZT12 n = 6, ZT16 =10, ZT20 n = 3; 12:12 25ºC: ZT0 n = 14, ZT4 n = 14, ZT8 n = 15, ZT12 n = 19, ZT16 =10, ZT20 n = 15; 12:12 10ºC: ZT0 n = 13, ZT4 n = 14, ZT8 n = 15, ZT12 n = 15, ZT16 =16, ZT20 n = 10.

To determine the effect of the photoperiod alone on PDF levels, the same experiment was conducted in flies entrained to either a long photoperiod or a short photoperiod but at constant temperature 25°C (Figure 1D-E). No differences in the MESOR (p = 0.06), amplitude (p=0.9) or phase (p=0.72) were observed between these conditions respectively. Furthermore, PDF abundance in both LP (p = 0.005) and SP (p = 0.034) were found to be rhythmic (Figure 1F). To test the effect of temperature, flies were reared at 25°C or 10°C in 12:12 LD cycles (Figure 1G-H). We observed a reduction in the MESOR at 10°C compared to 25°C (MESOR 25°C = 2.95, MESOR 10°C = 2.16, p = 5×10^−6^; Figure 1I). Moreover, a change in the phase of the peak in PDF level was observed, with an estimated peak at ZT14 in 10°C and at ZT23 in 25°C (p = 9×10^−4^; Figure 1I).

As we observed low PDF levels in simulated winter, we hypothesized that a reduction in *pdf* mRNA could be the cause. To test this hypothesis, we conducted fluorescent *in situ* hybridization (FISH) to detect *pdf* transcripts in brains of flies expressing *CD8::GFP* in *pdf+* cells. We subjected these flies to either simulated summer or simulated winter. As *pdf* mRNA expression does not show daily oscillation^51^, only one timepoint (ZT16) was assessed in this experiment. A clear signal, confined to CD8::GFP positive cells, was observed (Figure 2A). This signal was absent in brains from *pdf-*null mutants (*pdf*^*01*^), confirming the specificity of FISH library (Figure S1). A reduction in fluorescent signal was observed in the LNvs somas under simulated winter compared to flies reared in simulated summer (Figure 2B). Quantification of the signal from different brains show a consistent reduction in the normalized fluorescent intensity in the CD8::GFP+ cells consistent with lower levels of *pdf* mRNA (t(13) = 3.267, p = 0.0061, Figure 2C).

**Figure 2:**
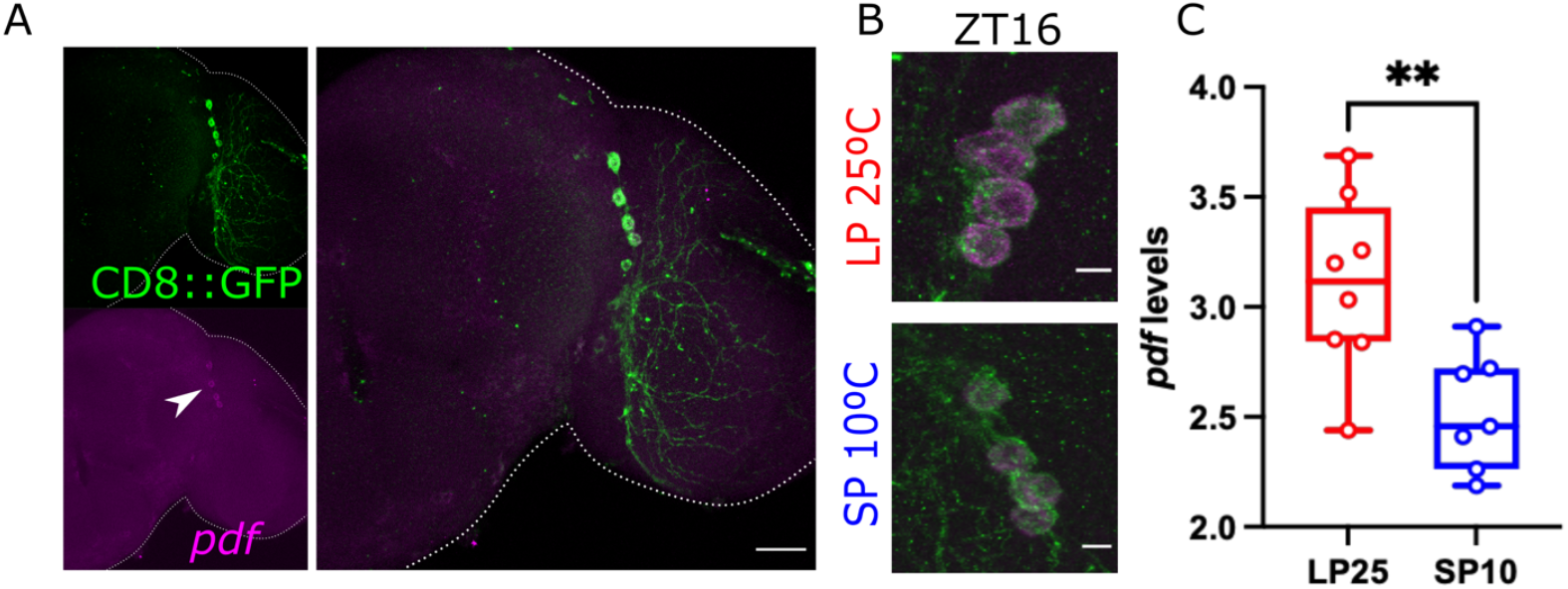
*pdf* mRNA is sensitive to seasonal cues. (A) *pdf* mRNA (magenta) was detected using fluorescence in situ hybridization (FISH) in whole brains of flies entrained for 7 days to simulated-summer (LP at 25ºC) or simulated-winter (SP at 10ºC). CD8::GFP was expressed using the *pdf-gal4* driver to detect the PDF-producing LNvs. (B) Representative images of LNvs of flies entrained to LP 25ºC (top panel) or SP 10ºC (bottom panel). (C) Quantification of the normalized intensity of the *pdf* signal at ZT16 is compared between conditions. Unpaired t-test, ** p <0.01. Number of brains imaged were LP25 n = 8, SP10 n = 7 brains. Scale bar in A is 25 μm and 5 μm in B.

### Adaptation of daily locomotor activity rhythm to cold temperature and short photoperiod depends on PDF

PDF is a modulator of daily locomotor activity in *Drosophila. pdf*^*01*^ mutants have previously been shown to exhibit an advancement in the evening peak of activity in LD and loss of rhythmicity under constant darkness^44^. As we observed a decrease in *pdf* mRNA and PDF protein in simulated winter or when flies are exposed to lower temperature, we reasoned that changes in *pdf* abundance should translate into changes in locomotor activity in response to photoperiodic and/or temperature changes, which has been shown to change over the calendar year in insects including flies. This would represent an effective readout of overall PDF abundance in flies. We therefore monitored daily activity in *pdf*^*01*^ flies compared to *pdf*^*01*^ mutants bearing a genomic rescue *pdf* transgene (*yw* ; *pdf{WT}* ; *pdf*^*01*^, hereafter denoted as genomic rescue) in selected conditions. First, flies were monitored in 12:12 LD cycles but exposed to either 25°C or 10°C. As previously described, *pdf*^*01*^ flies display an advance in evening activity peak at 25°C consistent across the 5 days of monitoring. This is not observed in the genomic rescue, which has normal timing of evening peak immediately before the lights off transition at dusk (Figure 3A-B). In contrast, at 10°C, an advancement in the evening peak was now observed even in the genomic rescue flies as they behaved similar to *pdf*^*01*^ mutants (Figure 3C-D). Then, to test the effect of photoperiod, we monitored locomotor activity in short day conditions (8:16 LD) at 25°C in *pdf*^*01*^ mutants and genomic rescue flies. Consistent with previous studies, *pdf*^*01*^ mutants showed a peak that closely follow the lights-off transition (Figure 3E-F).

**Figure 3.**
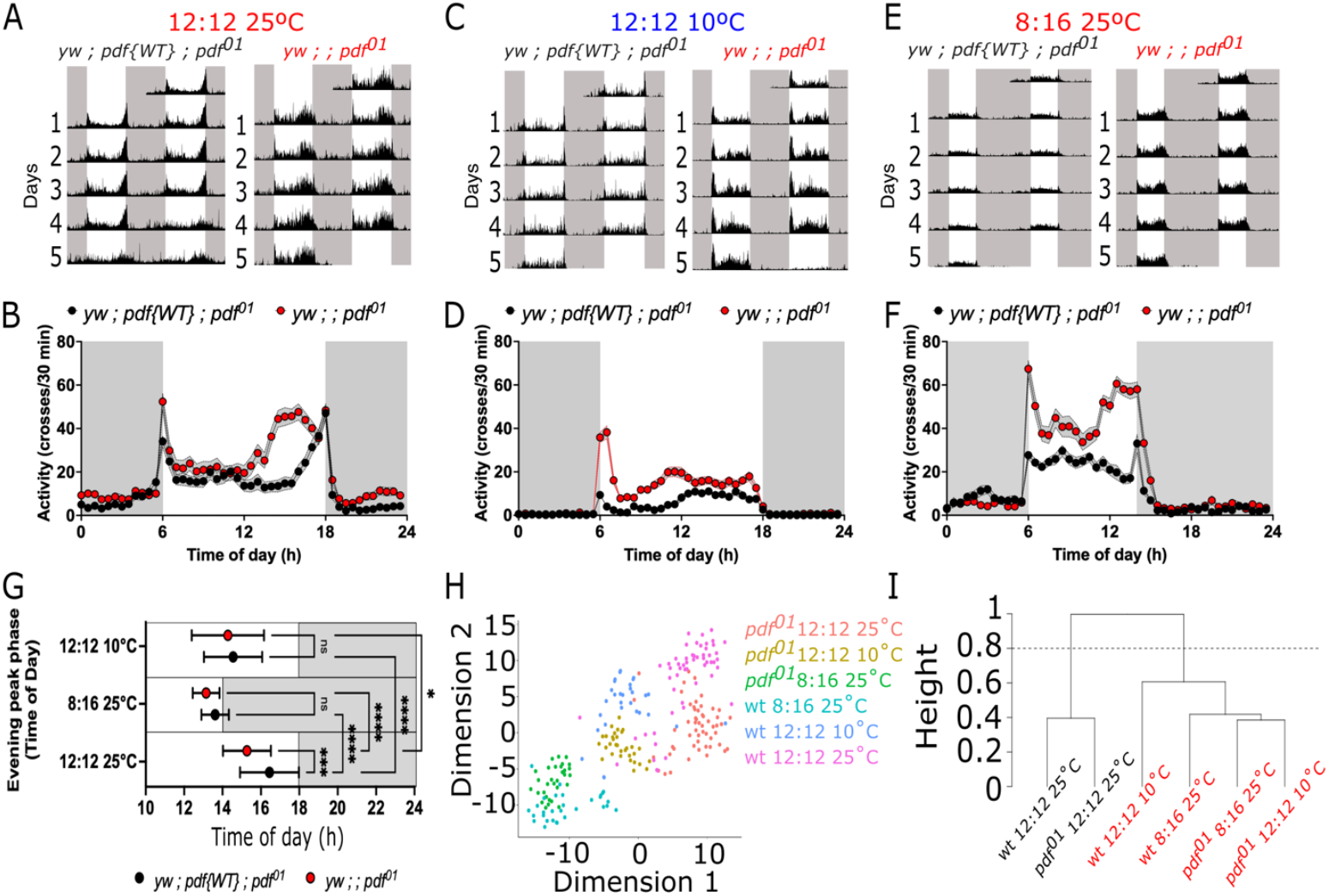
Locomotor adaptation to photoperiod and temperature in *pdf*-null mutants. Locomotor activity was monitored in *pdf*-null mutants (*yw* ; *pdf*^*01*^) and in flies expressing a wild-type copy of *pdf* (genomic rescue, *yw* ; *pdf{WT}* ; *pdf*^*01*^) subjected to (A-B) 12:12 LD at 25ºC, (C-D) 12:12 LD at 10ºC, or (E-F) 8:16 LD at 25ºC. (G) Comparison of the evening peak phase between *pdf*^*01*^ (red boxes) and the genomic rescue (black boxes) in different environmental conditions. (H) Variance in locomotor activity under these environmental conditions as tSNE of the normalized locomotor activity. Each dot represents an individual fly. (I) Hierarchical clustering of the normalized locomotor activity. Cut of the tree at 0.8 generates two distinct groups denotated as black and red labels. White boxes represent lights-on while black boxes represent lights-off in panels A-F. Data were analyzed using a two-way ANOVA with Bonferroni’s multiple comparison test. Number of flies used were *yw* ; *pdf{WT}* ; *pdf*^*01*^ 12:12 25°C n = 56, *yw* ; *pdf{WT}* ; *pdf*^*01*^ 8:16 25°C n = 30, *yw* ; *pdf{WT}* ; *pdf*^*01*^ 12:12 10°C n = 30, *yw* ; ; *pdf*^*01*^ 12:12 25°C n = 55, *yw* ; ; *pdf*^*01*^ 8:16 25°C n = 31, *yw* ; ; *pdf*^*01*^ 12:12 10°C n = 32.

As depicted in the eduction graphs (Figure 3B,D and F), the main differences on activity rhythm in LD are located in the timing of the evening peak. Thus, we compared the phase of the evening peak between the rescue and the mutant in all these conditions. *pdf*^*01*^ mutants have an advancement in the evening peak of activity compared to rescue flies in 12:12 LD at 25°C (F(2, 228) = 11.97, p = 0.0006). Changing either the temperature or photoperiod caused a phase shift in evening activity peak to earlier times of the day in the genomic rescue and *pdf*^*01*^ mutants (F(2, 228) = 69.61, p < 0.0001). The differences observed between rescue and *pdf*^*01*^ mutants are no longer observed at 12:12 10°C or 8:16 LD 25°C (Figure 3G).

When exposed to cold temperature, genomic rescue flies seem to behave like the *pdf*^*01*^ mutants at 25°C and 10°C (Figure 3A-D). To better visualize the overall similarities of fly behavior between the different datasets, we conducted tSNE analysis on the observed locomotor activity rhythms. We took the average daily activity profiles in LD for each fly and normalized the activity to the maximum activity in each fly. This enabled the comparison of the overall shape/structure of the activity of the normalized locomotor activity (Figure 3H). Overall, three distinct groups can be defined by the environmental conditions tested. While data from the genomic rescue and the *pdf*^*01*^ mutants at 25°C (magenta and red dots, respectively) cluster within two broad individual groups, data between the two genotypes at 10°C (blue and yellow dots) and 8:16 (green and cyan dots) are less differentiated. To further define these clustering, we used an unsupervised analysis of the normalized locomotor activity per group using hierarchical clustering (Figure 3I). Two main branches are observed, one with rescue and mutant flies at 12:12 at 25°C (black labels) and the other one containing both mutants and rescue flies at 8:16 and 10°C (red labels). These suggest that at these environmental conditions, the structure of the locomotor activity in mutant and rescue flies are similar to each other than to the flies at 12:12 25°C. Overall, these data suggest that temperature and photoperiod differentially modulate the evening activity in *pdf*^*01*^ mutants compared to control flies.

### PDF modulates overall activity level through action in *eya+* cells but its effect on timing of evening peak is not dependent on EYA

Upon release, PDF acts through activation of the G-coupled PDF receptor (PDFR). Thus, we reasoned that if PDF signaling acts through EYA to regulate changes in daily activity rhythms in response to photoperiodic and temperature changes, cells that are important in this process would coexpress *eya* and *pdfr*. To explore this, we first used a single-cell transcriptomic datasets available from the Fly Cell Atlas initiative^53^ to determine if *pdfr* is coexpressed with *eya* in the fly brain. Using SCOPE, a visualization tool for large sequencing dataset^54^, we differentiated all cell/tissue types in the scRNA-seq dataset based on overall expression (Figure S2A), and also colored them specifically by *pdfr* and *eya* expression (Figure S2B; green and red dots, respectively). Coexpression of both transcripts is observed in head tissue, gut, and testis cluster. Further examination showed that many of the cells expressing *eya* within the head cluster also express *pdfr* (Figure S2B; insert).

To determine if PDF-dependent changes in daily activity rhythms in response to photoperiodic and temperature as observed in Figure 3 is mediated via EYA, we monitored locomotor activity under 12:12 LD cycles at 25°C in flies where *pdfr* was knocked down in *eya+* cells using an *eya-Gal4* driver (Figure S3A)^24^. Interestingly, we did not observe a dramatic shift in the timing of the evening activity peak as in *pdf*^*01*^ flies. This suggests that PDF signaling alters timing of evening peak through mechanisms other than EYA. We did observe, however, an increase in the strength of the evening anticipation in *eya > pdfr-RNAi* flies compared to controls (Figure S3A-B). Notably the opposite is observed in flies driving a tethered version of PDF (t-PDF) using the same driver to activate PDF signaling (Figure S3C-D). This difference is even more pronounced when looking the overall activity level of flies in constant darkness. Knocking down *pdfr* in *eya+* cells results in an overall reduction in total locomotion (H(2, 86) = 17.89, p = 0.0001; Figure 3A-B), while activating PDF signaling by expressing t-PDF in *eya+* cells promotes an increase in activity (U = 333, p = 0.0384; Figure 3C-D). Similar to our observations in LD (Figure S3), no change in the phase of evening activity was observed in DD (Figure 3A-D).

Previous studies have shown that PDF modulates evening anticipation by its action on non-*pdf*-expressing clock cells within the CCNN^55^. Thus, it is possible that the effect observed while knocking down *pdfr* or expressing t-PDF in *eya+* cells is due to the modulation of these neurons. To test if *eya* is expressed in clock neurons, we detected EYA using a monoclonal antibody (Figure 4F) in flies carrying a endogenous GFP-tagged version of PER^56^ to mark neurons in CCNN (Figure 4E). EYA was detected in some PER-expressing cells including DN1, LNd and DN2 neurons (Figure 4G) consistent with a role of these clusters in locomotor activity.

**Figure 4.**
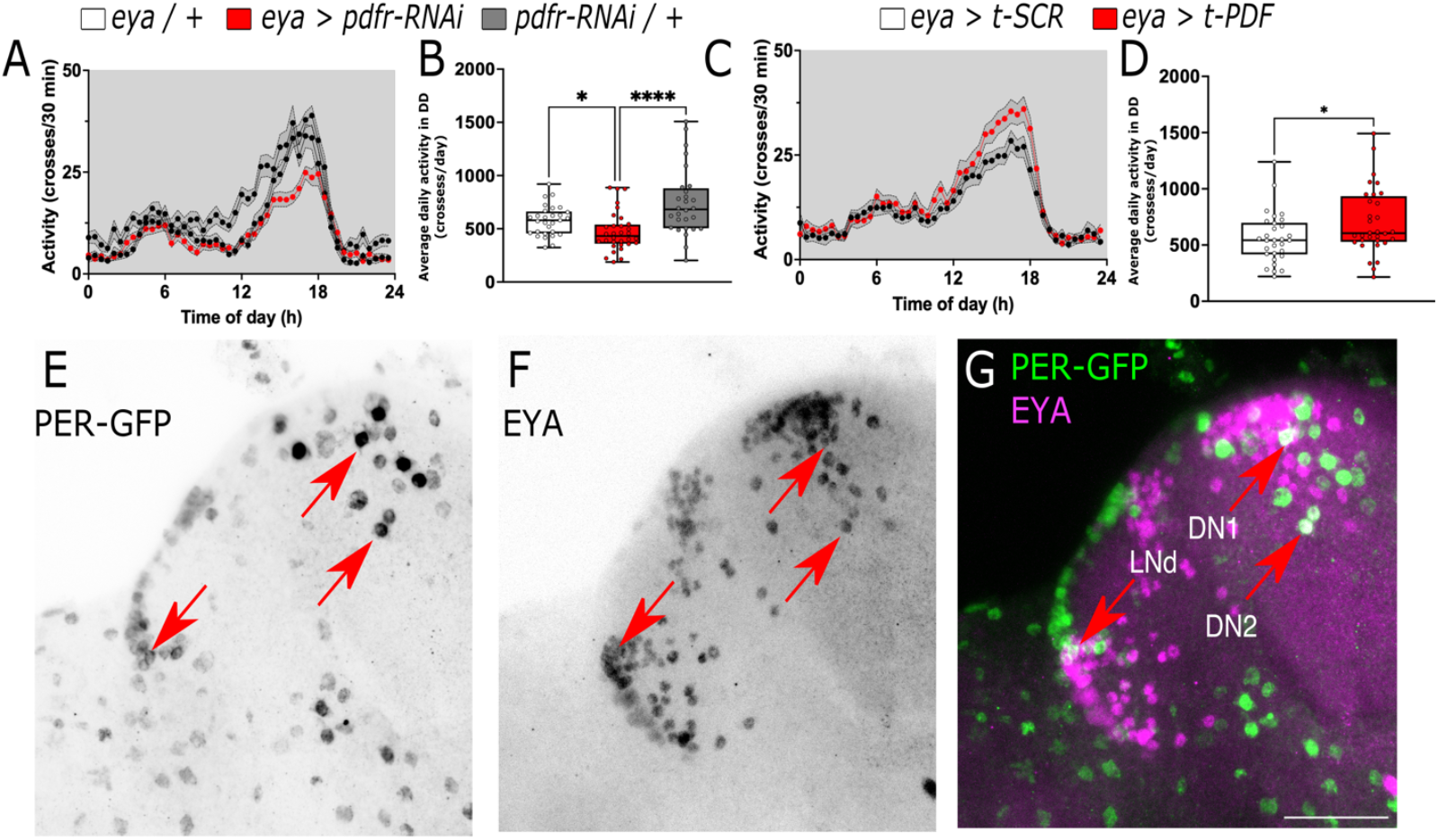
PDF modulates overall locomotion through *eya+* cells. (A-B) Average locomotor activity of flies in constant darkness at 25ºC where the PDF receptor was knocked down in *eya*+ cells (*eya > pdfr-RNAi* red line and box) compared to controls (*eya* / + and *pdfr-RNAi* / +, black and gray lines, respectively). (C-D) Average locomotor activity of flies where a membrane tethered version of PDF was expressed in *eya*+ cells (*eya > t-PDF* red line and box) compared to the expression of a scrambled version of the peptide (*eya > t-SCR* black line and box). (E-G) Confocal images of fly brain expressing endogenous GFP tagged PER (green) and stained against EYA (magenta). Red arrows points DN1, DN2 and LNd clock populations. Scale bar is 25 μm. Data in B were analyzed with Kruskal-Wallis test followed by Dunnett’s multiple comparison test, Mann-Whitney in D. Number of flies used were *eya / +* n = 29, *eya > pdfr-RNAi* n = 32, *pdfr-RNAi / +* n = 28, *eya > t-SCR* n = 31, *eya > t-PDF* n = 31, *eya / +* n = 30.

### PDF modulates ovary size through action in *eya+* insulin-producing cells

Expression of *eya* in the IPCs is critical for the regulation of seasonal reproductive dormancy^24^. To determine if PDF signaling works through EYA in IPCs to regulate seasonal phenotypes, we first tested if *pdfr* is coexpressed with *eya* in these cells. Similar to a previous report^57^, we observed *pdfr*+ cells widespread across the brain (Figure S2). A cluster of cells is observed in the PI where the IPCs are located (Figure 5A, inset). We used SCOPE to plot all the cells from the dilp2IPC SMART-seq dataset, generated from isolated IPCs by fluorescence-activated cell sorting^53^, and colored them by *pdfr* and *eya* (Figure 5B; green and red dots). We observed *eya* expression in a small number of cells and some of them also express *pdfr* (Figure 5B; yellow dots). We then detected EYA using a monoclonal antibody in *pdfr > CD8::GFP* brains, focusing on this cluster. As previously described^24^, EYA expression is observed within these cells, consistent with its expression in the IPCs, and is within the *pdfr+* cells (Figure 5C, blue arrows). Thus, both transcriptomic and immunohistochemistry data support the notion that EYA and PDFR are coexpressed in the IPCs in fly brains.

**Figure 5.**
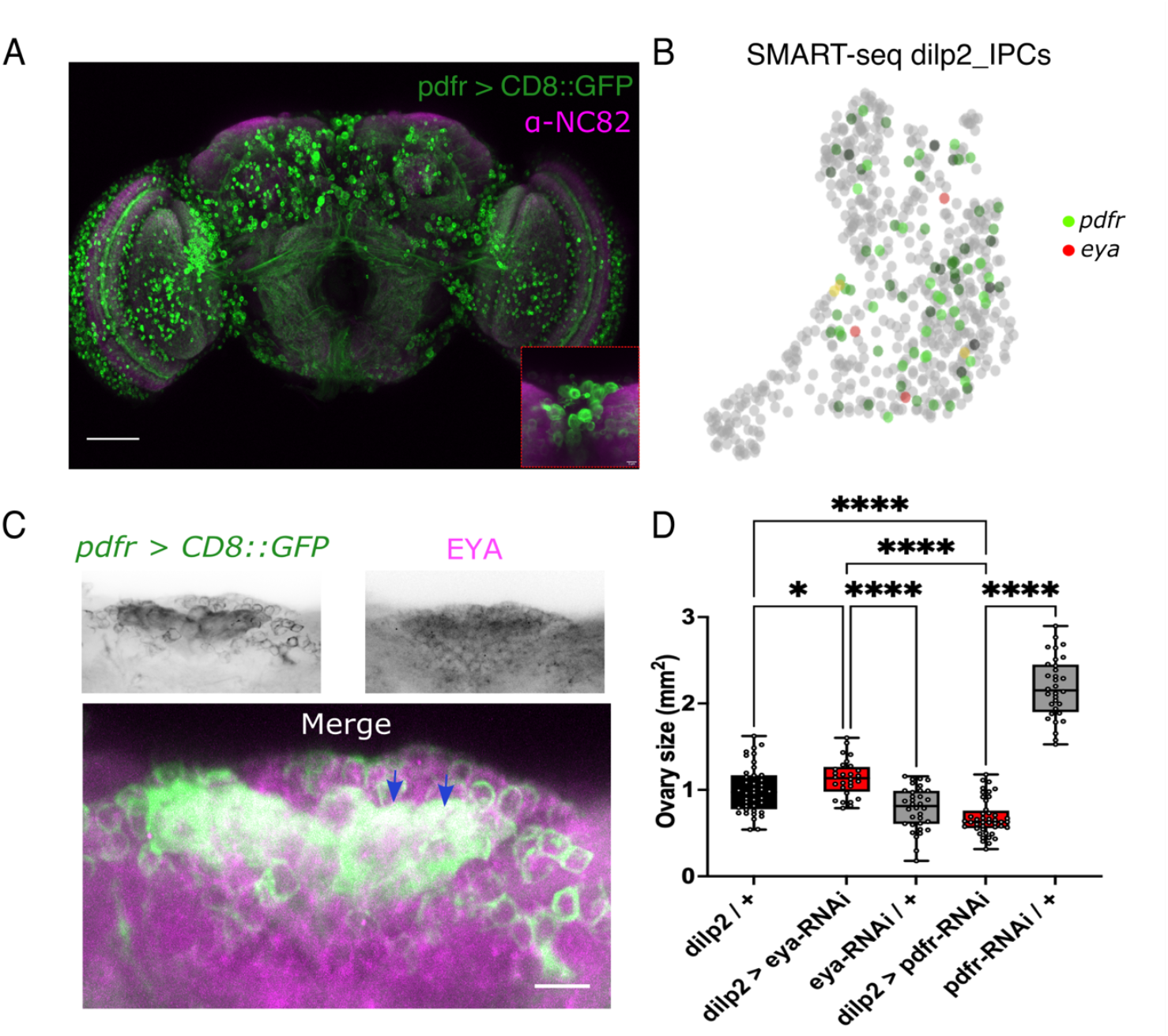
PDF and EYA have opposite effects in the IPCs to modulate ovary size under summer-like conditions (16:8 LD at 25°C) (A) Expression of CD8::GFP was driven in the *pdfr+* cells using a *pdfr-gal4* driver. Nc82 antibody was used as counterstaining. Expression of CD8::GFP is observed in several segments of the brain, including the *pars intercerebralis*, containing the IPCs (inset). Scale bar is 50μM. (B) SMART-seq dataset from *dilp2+* isolated IPCs from Fly Cell Atlas^53^ as UMPA, colored by *pdfr* (green) and *eya* (red) expression. Co-expression is shown as yellow. (C) EYA staining (upper right panel) in brains of flies expressing CD8::GFP in the *pdfr+* cells (*pdfr > CD8::GFP*, upper left panel). Merged image is observed in the bottom panel. Scale is 5 μm. (D) Comparison of ovary size in mm^2^ while knocking down *eya* (*dilp2 > eya-RNAi*) or knocking down *pdfr* (*eya > pdfr-RNAi*) compared to genetic controls (black and grey boxes). Data was analyzed using a one-way ANOVA followed by Holm-Šĺdák’s multiple comparison test. Number of flies used were *dilp2 / +* n = 40, *dilp2 > eya-RNAi* n = 28, *eya-RNAi / +* n = 34, *dilp2 > pdfr*-RNAi n = 45, *UAS-pdfr-RNAi / +* n = 32.

We showed that PDF levels decrease in winter-like conditions (Figures 1-2), while our previous study showed that EYA levels increase under similar conditions. This suggests opposite functions of PDF vs EYA. To validate the antagonistic relationship between PDF and EYA, we knocked down *pdfr* or *eya* in the IPCs and examined ovary size in summer-like conditions. We hypothesized that, knocking down *pdfr* in this condition, should render smaller ovaries and knocking down *eya* even in summer-like condition may produce bigger, more developed ovaries, as *eya* promotes winter physiology and reproductive dormancy. As expected, knocking down *pdfr* generated a reduction in the ovary size in summer conditions compared to the parental controls, while the same manipulation for *eya* produced the opposite effect, suggesting opposite roles of EYA and PDF in ovary development in summer-like conditions (F(4, 173) = 177.8, p < 0.0001; Figure 5D). We did not perform a parallel experiment in winter-like conditions given we have previously shown that knocking down *eya* in winter-like conditions resulted in lower levels of reproductive dormancy^24^. Moreover, we do not anticipate further reduction of ovary size if we knock down *pdfr* in winter-like conditions when ovaries are already undeveloped with no eggs. Taken together, these data suggest that PDF modulates ovary size through its action in *eya* expressing cells and PDF and EYA function antagonistically.

### EYA protein but not mRNA is modulated by PDF

We next sought to determine if PDF could act upstream of EYA. First, we quantified *eya* mRNA in heads of *pdf*^*01*^ and genomic rescue flies entrained in 12:12 LD cycles at 25°C. We observed no differences in *eya* abundance at all time points over the 24 hour cycle (F(1, 56) = 0.9281, p = 0.3395; Figure 6A). Nonetheless, statistical analysis of rhythmicity revealed that *eya* exhibits rhythmic expression in the genomic rescue flies (RAIN p = 0.0078), while it is not rhythmic in *pdf*^*01*^ mutants (RAIN p = 0.833). Then, to determine if EYA protein level is different between the two genotypes, we assayed EYA abundance in protein extracts of heads from *pdf*^*01*^ mutants or genomic rescue flies. Remarkably, we observed a dramatic increase in the overall amount of EYA protein in *pdf*^*01*^ mutants as compared to the genomic rescue throughout the 24 hour cycle (F(1, 55) = 50.01, p < 0.0001; Figure 6B-C). Similar to our observation with *eya* mRNA, EYA protein showed daily rhythmic expression only in the genomic rescue (RAIN p = 0.0398), no robust daily rhythmicity was observed in *pdf*^*01*^ mutants (RAIN p = 0.667).

**Figure 6.**
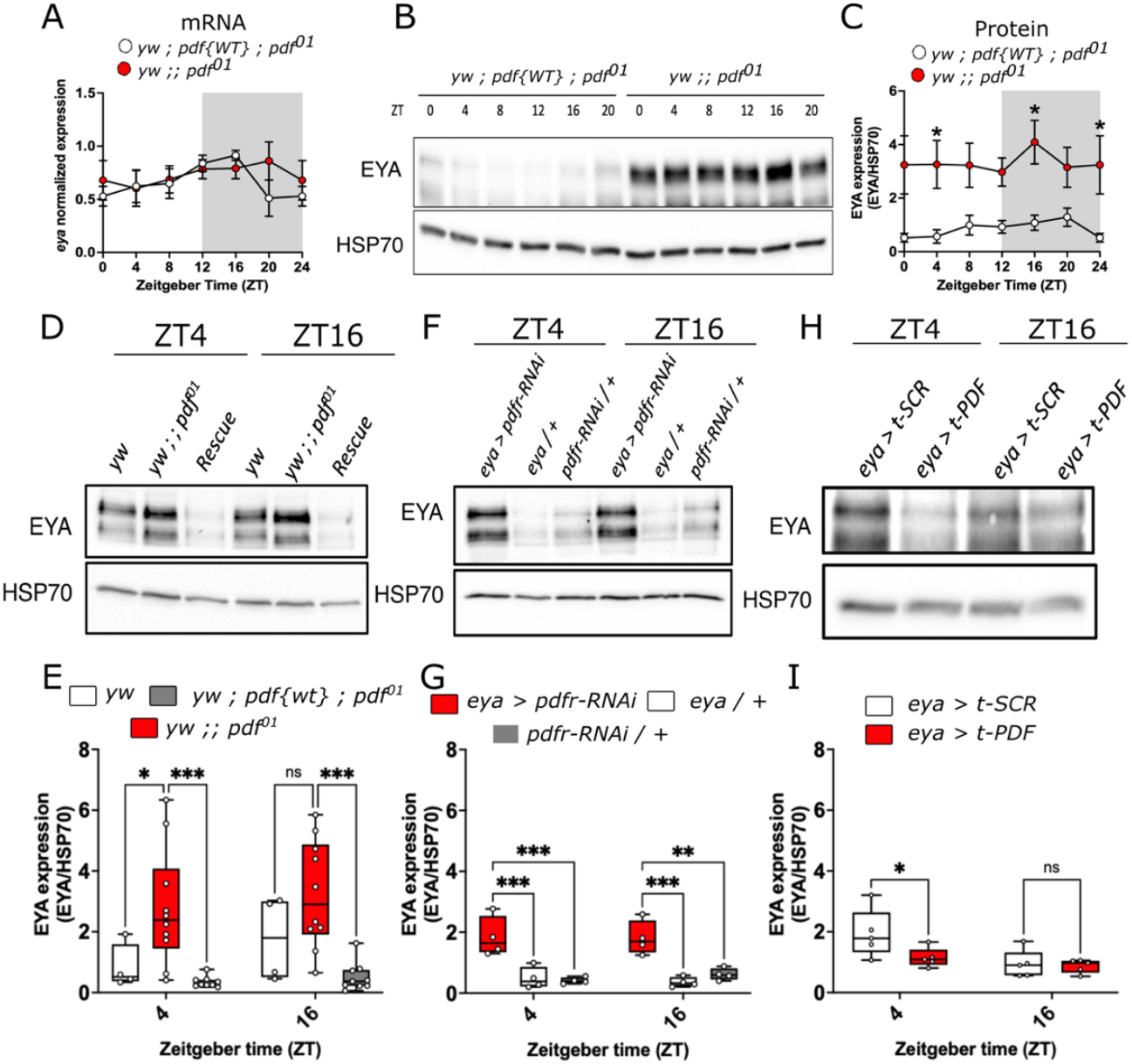
PDF modulates EYA protein but not *eya* mRNA expression. (A) *eya* mRNA daily oscillations in *pdf*-null mutants (*yw* ; ; *pdf*^*01*^ flies, red dots, RAIN p = 0.833) and in flies expressing a *pdf* genomic rescue (*yw* ; *pdf{WT}* ; *pdf*^*01*^, white dots, RAIN p = 0.0078) in 12:12 LD cycles at 25ºC. (B) Representative western blot and (C) quantification of EYA levels in whole-head lysates from *pdf* null-mutant (red dots, RAIN p = 0.677) and in the genomic rescue (white dots, RAIN p = 0.039) at ZT0, 4, 8, 12, 16 and 20 in 12:12 LD 25ºC cycles. (D) Representative western blot and (E) quantification of EYA levels from whole head lysates of *yw* (genetic control, white boxes), *pdf-null* mutants (red boxes) and the genomic rescue (grey boxes) at ZT4 and ZT16 in 12:12 LD 25ºC condition. (F) Representative western blot and (G) quantification of EYA levels in head lysates of flies where the PDF receptor was knocked down in *eya+* cells (*eya > pdfr-RNAi*, red boxes) and controls (*eya* / + and *pdfr-RNAi* / +, white and grey boxes, respectively) at ZT4 and ZT16. (H) Representative western blot and (I) quantification of EYA levels in flies expressing a membrane-tethered version of PDF in *eya+* cells (*eya > t-PDF*, red boxes) compared to a scrambled peptide as control (*eya > t-SCR*, white boxes) at ZT4 and ZT16. Data were analyzed using a two-way ANOVA followed by Tukey’s post hoc test in B and C, and Šĺdák’s multiple comparison test in E, G and I. HSP70 was used for normalization in all cases. n > 3 biological replicates, each consisting of around 50 flies for each genotype.

To confirm these findings, we repeated the experiment, this time with a wild type (WT) control fly in the same genetic background (*yw*), given that the genomic rescue might not fully recapitulate the WT phenotype. An increase in EYA levels was still observed at ZT4 in *pdf*^*01*^ mutants compared to control WT flies or the genomic rescue (F(2, 40) = 17.92, p < 0.0001; Figure 6D-E). To test if the effect of *pdf*^*01*^ mutation in EYA protein levels is mediated by function of PDF signaling in *eya+* cells, we knocked down *pdfr* using the *eya-Gal4* driver. Similarly to the *pdf*^*01*^ mutant, an increase in the levels of EYA was observed in these flies (F(2, 18) = 30.64, p < 0.0001; Figure 6F-G). Conversely, expressing t-PDF to activate PDF signaling in the same *eya+* cells triggered the opposite effect; flies expressing t-PDF showed lower EYA levels than flies expressing a scrambled (SCR) peptide (F(1, 16) = 8.268, p = 0.0110; Figure 6H-I). These results suggest that PDF signaling can modulate EYA protein levels.

### PKA phosphorylates EYA to modulate its stability

Because PDF negatively affected EYA protein level with no changes in overall *eya* mRNA (Figure 6), we hypothesized that PDF modulates EYA stability through post-translational modification. PDFR activation leads to increased cAMP^40,58^ and subsequent activation of Protein kinase A (PKA) (Figure 7A). Interestingly, we found using pkaPS^59^ that EYA contains seven potential PKA phosphorylation sites: three of them in the C-terminal, one between the second Proline-Serine-Threonine (PST) rich domain and the EYA domain (ED1), and two within the ED1 domain. To test if PKA could phosphorylate EYA, we expressed EYA alone in *Drosophila* S2 cells or together with PKA-C1, the enzymatically active subunit of this kinase^60^. We observed slower migrating EYA isoforms when EYA and PKA-C1 are coexpressed (Figure 7B). When treated with lambda phosphatase (λPP), we observed a clear mobility shift to faster migrating isoforms indicating that EYA is indeed phosphorylated by PKA-C1 in S2 cells. This data combined with the observation of EYA accumulation upon knockdown of *pdfr* (Figure 6) suggest that PKA potentially phosphorylates EYA to modulate its stability. To test this, we conducted a cycloheximide (CHX) chase experiment in S2 cells transfected with EYA alone or co-transfected with EYA and PKA-C1. Samples were collected at 0, 3, 6 and 9 hours after CHX application (Figure 7C). PKA expression increased EYA degradation rate, evidenced by a significant decrease in EYA levels after 3 hours of CHX addition as compared to the condition without PKA (F(1, 24) = 7.689, p = 0.0106; Figure 7C-D). Moreover, a mobility shift to higher molecular weight isoforms in EYA is clearly observed at 3, 6 and 9 hours post CHX in the presence of PKA-C1 coexpression (Figure 7C). Overall, our results suggest that PDF promotes PKA-dependent EYA phosphorylation and degradation.

**Figure 7.**
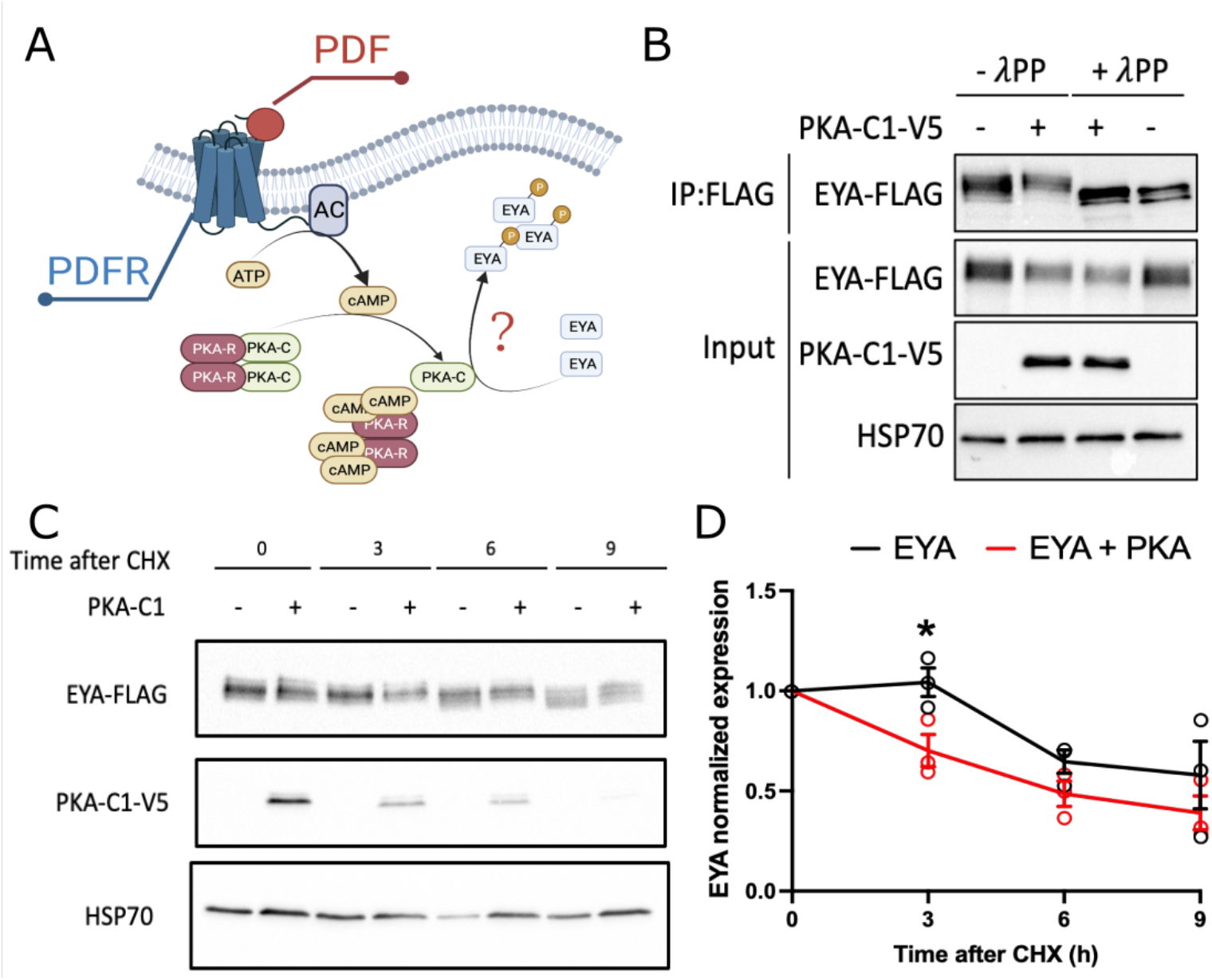
PKA phosphorylates EYA and mediates its degradation. (A) PDF receptor triggers the activation of adenylate cyclase (AC) which produces cAMP. The binding of cAMP to the regulatory subunits of the protein kinase A (PKA-R) releases the catalytic subunits of the kinase (PKA-C) permitting their action. EYA has potential sites for PKA phosphorylation. (B) *Drosophila* S2 cells co-expressing EYA-FLAG and the active C1 domain of PKA tagged with V5 (PKA-C1-V5 + condition) or EYA alone. Samples were then subjected to phosphatase treatment (+λPP) and compared to samples that were mock treated (-λPP). Immunoprecipitated EYA was detected using α-FLAG in the input. α-V5 and α-HSP70 were used to detect PKA-C1 and HSP70 (n>3, representative image is shown). (C) Representative western blot of a CHX chase assay in S2 cells co-expressing EYA-FLAG alone or with PKA-C1-V5. (D) Quantification showing the rates of EYA degradation in absence (black line) or presence (red line) of PKA-C1. Data analyzed with two-way ANOVA with Holm-Šĺdák’s multiple comparison test. Each dot represents an independent experiment (n = 3, * p < 0.05).

## Discussion

The mechanisms by which organisms sense photoperiod and temperature to adapt to seasons have been studied for many years, but the core molecular components involved in this process are still unclear. In this study we showed that the circadian neuropeptide PDF can respond to environmental cues to modulate seasonal phenotypes via EYA-dependent and independent mechanisms (Figure 8). Our findings suggest that PDF level is responsive to both photoperiodic and temperature changes. We proposed that under summer days, high PDF levels promote EYA degradation via PKA-dependent phosphorylation. In contrast, under winter days, a reduction in PDF drives accumulation of EYA to induce EYA-dependent changes in seasonal phenotypes, such as reproductive dormancy and changes in overall activity levels.

**Figure 8.**
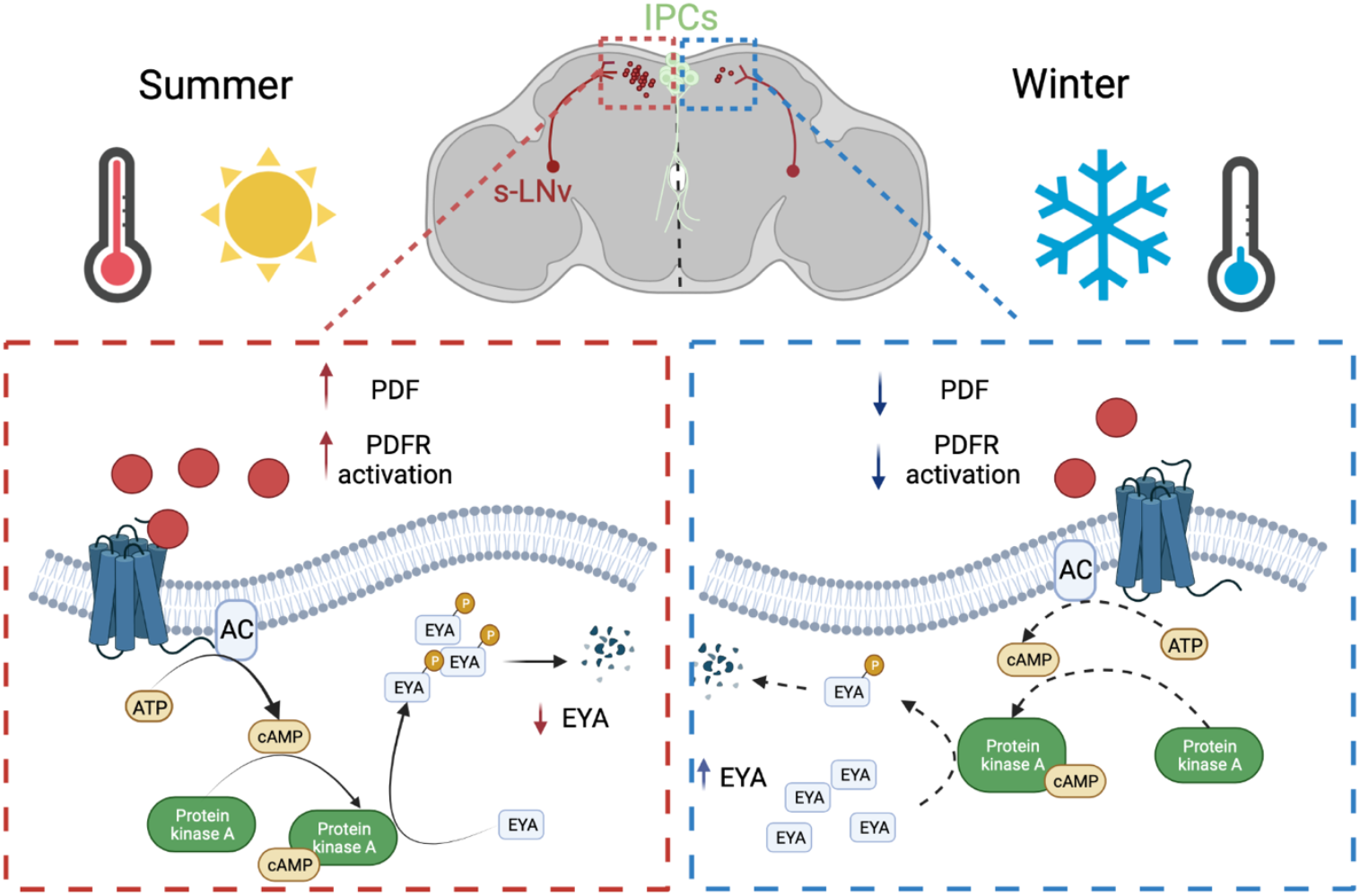
Model on PDF/PDFR contribution to seasonal physiology through EYA expressed in the Insulin-Producing Cells (IPCs). During summer, upon long days and high temperatures, PDF expression is increased potentially generating more PDFR activation and increase in active PKA. This in turn promotes EYA phosphorylation and degradation. During winter, with short days and low temperatures, a decline in PDF expression would promote EYA accumulation by a reduction of its PKA-mediated phosphorylation.

While other studies have explored the differences in PDF levels with environmental conditions, the results have been ambiguous^61,62^. Here we showed that during short and cold days, PDF levels are greatly reduced compared to long and warm days, effect driven by a reduction in *pdf* mRNA levels. This is consistent with a previous observation from our lab showing *pdf* as one of the transcripts that is downregulated in head of *Drosophila suzukii* winter morphs^63^. Overall changes in PDF can be largely explained by temperature, however, it is not clear how temperature changes results in changes in *pdf* levels. Several factors have been found to modulate *pdf*/PDF expression. For instance, neuronal activity and expression of CLK repress *pdf* transcription, possibly through Hormone Receptor 38 (HR38) and stripe (SR)^37^. *pdf* levels have also been shown to increase in response to high-intensity light through HR38 action^52^. In contrast, expression of VRILLE, a repressor in a secondary TTFL, promotes *pdf*/PDF mRNA and protein accumulation in the sLNvs through a post-transcriptional mechanism^38^. Hence, it is possible that changes in core clock components are involved in the reduction in *pdf*/PDF levels with cold temperatures. Interestingly, some core clock components are sensitive to temperature^64–67^. Alternative splicing of *per* and *tim* occurs at low temperature^64–68^. At 18°C, there is an increase in splicing of an intron located in the 3’ end of *per*, which leads to a phase advance in *per* levels, which ultimately drives a phase advance in the evening peak of locomotor activtiy^65^. We showed that at 10°C, control flies also display an advancement in the evening peak. This locomotor activity pattern resembled that of *pdf*^*01*^ flies at 25°C, closely matching the lights off transition of short days. Thus, it is possible that these changes rely on a *per/pdf* dependent pathway. At similar temperature, an intron retention event in *tim*, modulated by P-element somatic inhibitor, generates a shorter version of *tim* named *short and cold* (*tim-sc*)^67,68^. Interestingly, overexpression of *tim-sc* using a *tim-Gal4* driver generates akin but less pronounced changes in evening anticipation with flies initiating evening anticipation at around ZT7, similar to the time it starts in *pdf*^*01*^ at 25°C or genomic rescue flies at 10°C^67^. We previously showed that TIM-SC protein is indeed expressed at 10°C and experiments in S2 cells showed that TIM-SC interacts with EYA to modulate its stability^24^. However, it is not known if this isoform has any impact on the function of the molecular clock, ultimately affecting PDF levels. Martin Anduaga et al.^67^ showed that TIM-SC apparently has reduced binding to PER compared to TIM-L. Thus, the presence of TIM-SC during cold temperatures could potentially reduce PER stability or its translocation to the nucleus, causing CLK-CYC to promote the *pdf* reduction through HR38 and SR.

Considering our hypothesis that PDF signaling conveys photoperiodic and temperature cues to regulate seasonality via EYA and the previously identified role of EYA in winter physiology^17,24,33^, we expect that knocking down *pdfr* or activating PDF signaling by overexpressing t-PDF in *eya+* cells would alter activity level as a signal of seasonal adaptation. Indeed, knocking down *pdfr* decreased overall activity akin to a winter-like quiescent state while expressing *t-pdf* in the same cells rendered the opposite effect. This is consistent with a role of PDF in promoting activity in the summer. Using immunohistochemistry, we showed that EYA is expressed in PER-expressing neurons in the CCNN. Based on their anatomical location, we proposed they correspond to DN1, DN2 and LNds clusters^69^. Both, DN1, in particular the posterior cluster DN1p, and the LNds have been shown to modulate the evening anticipatory activity downstream of the s-LNvs^52,70,71^. Interestingly, the effect of the DN1ps and LNds seems to be opposite. While PDF inhibits the DN1ps in high-intensity light, it promotes LNds effect on evening anticipation^52,71,72^. Under our experimental conditions, the effect observed by modulating PDF is most likely mediated by LNds, although further experiments would be needed to assert this conclusion. The DN1ps also modulate the effects of temperature on daily locomotion^72–74^. The activity of the DN1ps is temperature dependent and they modulate sleep and locomotor activity in a temperature dependent manner^72,74^. It is thus possible that under winter-like conditions, DN1ps lead the phase advancement of evening anticipation, which allow flies to be exposed to light, necessary to keep them warm during winter-like days. Winter days also provide low intensity light that would potentially weaken the control of PDF over these neurons.

Once the temperature and photoperiod information are processed by the CCNN, the cues are then conveyed to circadian output circuits like the *pars intercerebralis* to modulate seasonal adaptations, including locomotion and reproductive status. In concordance with our hypothesis, it was shown that DN1s convey information from s-LNvs to modulate evening anticipation by acting in PI neurons expressing the neuropeptide diuretic hormone 44 (DH44). DH44 in turn modulates *hugin+* neurons that project to the ventral nerve cord to modulate locomotion^35,75^. In parallel, the IPCs receive direct inputs from the s-LNvs to modulate seasonal reproduction. Activating PDF cells increases egg laying at 25ºC^76^, and the same manipulation at 12ºC inhibits reproductive dormancy^50^. In contrast, silencing the same neurons promotes this winter adaptation, likely through a direct effect of PDF on the IPCs. This is because *pdfr* in the IPCs is sufficient to rescue dormancy levels in a *pdfr* mutant^50^. Consistent with these data, we showed that knocking down *pdfr* in the IPCs reduced ovary size in flies reared under long and warm days. Taken together, these data suggest that PDF functions in the IPCs to promote summer physiology.

Our previous findings showed that EYA promotes winter physiology. Increased *eya* expression in IPCs is sufficient to trigger dormancy while knocking it down caused increased ovary size^24^. We were able to replicate our previous results and observed that knocking down *eya* results in bigger ovaries even in long and warm days. Notably, this is opposite to the phenotype observed when knocking down *pdfr*, suggesting that PDF signaling may be antagonizing EYA function. Furthermore, we showed that cold temperature and short days trigger a reduction in *pdf/*PDF mRNA and protein, the response being opposite of what we previously observed for EYA^24^. Because PDF and the s-LNvs are upstream the IPCs, we reasoned that PDF may be upstream and convey environmental signals to modulate EYA. Our transcriptomic and immunohistochemistry analysis showed that PDFR is coexpressed with EYA in the IPCs, consistent with previous findings showing expression of both proteins in these cells^24,77^. EYA levels are increased in *pdf*^*01*^ mutants, consistent with a role of PDF in inhibiting EYA expression. Additionally, expression of t-PDF in *eya*+ cells was sufficient to decrease EYA levels, while an increase in EYA was observed when *pdfr* was knocked down. This strongly suggest a direct inhibiting role of PDF on EYA. We suggest that the action of PDF signaling cascade would be involved in this. Consistent with this notion, we showed in S2 cells that the PKA-C1, the catalytic subunit of PKA and downstream target of PDF signaling, can modulate EYA levels by phosphorylation-dependent degradation. PKA-C1 is also constitutively expressed in the IPCs, thus a similar process is likely happening *in vivo*^78^.

Finally, a similar configuration could be driving seasonal adaptations in other animals. Several studies have suggested a functional homology between the vasoactive intestinal peptide (VIP) and PDF in circadian rhythms and seasonal adaptations^79,80^. VIP is expressed by neurons located in the suprachiasmatic nucleus (SCN), the main hub for circadian timekeeping in the brain^81^. This peptide is important for circadian synchronization of the SCN individual clocks and activates the VPAC2 receptor, a class B G-protein coupled receptor that increases cAMP/PKA upon activation, similar to PDFR in *Drosophila*^79^. VIP is also indispensable for seasonal adaptations, as VIP knockout mice do not display behavioral or physiological adaptations while entrain to different photoperiods^80^. VIP is also expressed in the hypothalamus in birds and its levels are coupled to photoperiod and hormonal adaptations^82,83^. It would be interesting to explore possible modulations of EYA by VIP in the *pars tuberalis* and its effect on seasonal adaptations.

In summary, our study proposed a mechanism by which the circadian clock conveys photoperiod and temperature to clock outputs, to modulate seasonal adaptations. While we attribute the connection between the clock and the outputs to a role of a circadian neuropeptide, the molecular mechanism driving the modulation of the neuropeptide in response to environmental cues are still an open question. Furthermore, our study provides additional support for a role of EYA in seasonal adaptations and uncovers the importance of EYA post-translational modifications in the regulation of the seasonal biology.

## Materials and Methods

### Fly stocks

Flies were reared on standard corn-yeast diet and kept at 25ºC on a 12 hour light (L): 12 hour dark (D) cycle (12:12 LD). Fly stock were either obtained from Bloomington *Drosophila* Stock Center (BDSC), Vienna *Drosophila* Research Center (VDRC) or gifts from other labs.

Immunofluorescence was conducted in whole-mount brain of flies constitutively expressing the membrane-bound GFP under the *pdf-Gal4* driver control (*pdf-Gal4; UAS-CD8::GFP*, kind gift from Dr. Yong Zhang). Flies expressing endogenous PER tagged with GFP (PER-AID-GFP, kind gift from Dr. Yong Zhang) were used for co-stainig with EYA in Figure 4.

The previously validated^24^ *eya*-Gal4 line (BDSC stock no. 49292) was used to target *eya* expressing cells in behavioral experiments and western blotting. The *pdfr* knockdown in *eya+* cells was achieved by crossing these flies with *UAS-pdfr-RNAi* (BDSC stock no. 38347, described in Herrero et al.^84^). The activation of PDFR was achieved by expressing a membrane tethered version of PDF (t-PDF, BDSC stock no. 81111, described in Choi et al.^85^). Parental flies crossed with *w*^*1118*^ were used as genetic controls for the knockdown while a scrambled version of *pdf* (t-SCR, BDSC stock no. 81113) crossed with *eya*-Gal4 was used as control for *eya-Gal4 > UAS-t-PDF* flies.

### Immunofluorescence

Newly emerged flies were collected and placed under 12:12 LD cycles at 25ºC for 3 days before switching them to long photoperiod (16:8 LD at 25ºC), short photoperiod (8:16 LD at 25ºC), low temperature (12:12 LD at 10ºC), simulated winter (8:16 LD at 10ºC), simulated summer (16:8 LD at 25ºC) or kept in equinox conditions (12:12 LD at 25ºC) for 7 days. Flies were collected at ZT0, 4, 8, 12, 16 and 20 hours where ZT0 represents lights on time (Zeitgeber Time; ZT), fixed for 40 minutes in 4% PFA and dissected immediately. Brains were re-fixed in 4% PFA for 20 minutes, washed 3 times in 0.2% PBST, 15 minutes per wash, and then incubated in blocking solution (5% NGS, 0.05% Sodium Azide in 0.2% PBST) for 30 minutes. After that, brains were incubated with primary antibody for 2 days at 4ºC. Brains were then washed 3 times in 0.2% PBST, 15 minutes per wash, and then incubated with a secondary antibody tagged with a particular fluorophore. A list of antibodies source and dilutions is provided in Table S1. Finally, brains were mounted using ProLong Diamond Antifade Mountant (Life Technologies, Carlsbad, CA) and imaged using a Leica SP5 confocal microscope equipped with excitation diodes at 638 nm and OPSL exciting at 488. For the experiments determining PDF levels in the s-LNvs dorsal terminals, images were taken every 2 μm using a 40x oil objective and digital zoom. For images of the whole brain, a 20x objective was used instead. All images were analyzed using Fiji and PDF levels were assessed as in Hidalgo et al.^86^ using an automated script.

### Fluorescence *in situ* hybridization (FISH)

FISH was carried out as described in Long et al.^87^, with minor modifications. Briefly, flies were fixed in 4% PFA for 40 minutes and brains were dissected in cold 0.5% PBST solution. Dissected brains were re-fixed in 2% PFA for 55 minutes, washed with 0.5% PBST and then permeated with 5% acetic acid for 5 min on ice. Autofluorescence quenching was achieved by incubating the brains with 1% sodium borohydride for 30 min. Sequences of *pdf* probes were the same as described in Long et al.^87^, tagged with CAL Fluor Red 635, purchased from Biosearch Technologies (Stellaris RNA FISH probes) (Petaluma, CA). Mounting was conducted as for immunofluorescence, using ProLong Diamond Antifade Mountant (Life Technologies).

### *Drosophila* Activity Monitoring (DAM) system

Locomotor activity was measured using the DAM system (TriKinetics, MA)^88^. Flies were anesthetized under CO_2_ and loaded in individual glass tubes containing fly food (5% sucrose and 2% Bacto agar in distilled water). Tubes were loaded in monitors and placed inside the incubator at 25ºC under 12:12 LD cycle. After 2 days of acclimation, locomotor activity was recorded for 5 days in LD conditions and released into constant darkness (DD) for 5 days. Only flies that survived until the last day of the experiments were considered for analysis. ZT0 was considered as the time in which the lights-ON transition occurs. Eduction graphs were generated for LD and DD for the 5 days in both conditions and the total locomotor activity was quantified as the sum of all the locomotor events in that given period. The evening peak amplitude was calculated as the sum of the activity between ZT9 and ZT12. Evening anticipation was calculated as the difference between the normalized averaged activity in ZT9–11.5 and ZT5–7.5, as previously described^86^.

### Ovary size quantification

Virgin females were placed in simulated summer conditions (25ºC, 16:8 LD) for 28 days and then dissected in 0.2% PBST (0.2% Triton X-100 in 1X PBS)^24^. Ovaries were imaged using a EVOS microscope (Life Technologies, CA) and ovary size was determined by measuring the surface area (mm^2^) using Fiji software.

### Total RNA extraction and quantitative RT-PCR

Flies were collected at the indicated timepoints on dry ice and then heads were collected and stored at -80°C until sample processing. Around 20-50 μL of heads per condition were processed using TRI reagent (Sigma-Aldrich, St. Louis, MO), following manufacturer’s instructions. Total RNA was quantified and 1 μg was used as starting material for reverse transcription using the Superscript IV cDNA synthesis kit (Life Technologies) following manufacturer’s instruction. Quantitative PCR was performed using SsoAdvanced Universal SYBR green super mix (Bio Rad, Hercules, CA), with the following the program: initial 95°C for 30 seconds, then 40 cycles of 95°C for 5 seconds, followed by an annealing/extension phase at 60°C for 30 seconds. Primers to detect *eya* were 5’-GAGGCCTGGCTACAGATACG-3’ and 5’-AGTTGCGTGGAGGTTACCAG-3’, while *cbp20* (5’-GTCTGATTCGTGTGGACTGG-3’ and 5’-CAACAGTTTGCCATAACCCC-3’) was used for normalization. Data were analyzed using the ΔΔCt method.

### Protein extraction from fly heads and western blotting

Protein extraction from fly heads was conducted as previously described^24,89–91^. Flies were collected at the indicated timepoints and flash frozen using dry ice. Heads were then collected and ground on protein extraction buffer (0.1% Glycerol, 20mM Hepes pH7.5, 50mM KCl, 2mM EDTA, 1% Triton X-100, 0.40% NP 40, 1mM DTT, 10μg/mL Aprotinin, 5μg/mL Leupeptin, 1μg/mL Pepstatin A and 0.5mM PMSF). Protein samples were kept at -80ºC in SDS loading buffer until electrophoresis in 8% SDS-PAGE gels. Transfer was conducted in a Trans-Blot semi-dry transfer system (Bio Rad) for 45 minutes to nitrocellulose membranes (0.45μm, Bio Rad). 5% blocking reagent (Bio Rad) in 0.05% TBST (0.05% Tween20 in 1XTBS) was used to block for 1 hour before antibody incubation overnight at 4ºC. Antibodies and dilutions are indicated in Table S1. Anti-mouse IgG linked to HRP (Cytiva Life Sciences, Malborough, MA) was used as a secondary antibody at a dilution of 1:1,000 incubated for 1 hour. Membranes were incubated for 5 min with Clarity reagent (Bio Rad) and imaged on ChemiDoc MP Imaging system. Image analysis was performed using Image J.

### *Drosophila* Schneider 2 (S2) cell culture experiments

*Drosophila* S2 cells were maintained at 22ºC in Schneider’s *Drosophila* medium (Life Technologies) supplemented with 10% Fetal Bovine Serum (VWR, Radnor, PA). For both cycloheximide (CHX) chase and phosphatase experiments, on day one, 3 × 10^6^ cells were transiently transfected with 0.8 μg *pAc-eya-3XFLAG-6XHis* and 0.2 μg *pMT-PKA-C1-V5*. PKA-C1 expression was induced 24 hours after transfection by addition of 500 μM CuSO_4_. For phosphatase treatment, cells were harvested 24 hours after kinase induction. Proteins were extracted using EB2 supplemented with 1X PhosSTOP (Roche, Indianapolis, IN) and FLAG immunoprecipitation was performed using anti-FLAG M2 agarose (Sigma) following protocol described in Lam et al.^90^ Immunocomplexes were then treated with Lambda phosphatase λ-PP or mock treated (New England Biolabs, Ipswich, MA) and then resolved in 8% SDS-PAGE. For cycloheximide (CHX) chase assay to measure protein degradation, CHX was added to the transfected cells, again at 24 hours after kinase induction to a final concentration of 10 μg/ml. S2 cells were harvested immediately, or at 3, 6, 9 hours after addition of CHX. Upon cell harvest, proteins were extracted using EB2 (20 mM HEPES pH 7.5, 100 mM KCl, 5% glycerol, 5 mM EDTA, 1 mM DTT, 0.1% Triton X-100, 10 μg/ml Aprotinin, 5 μg/ml Leupeptin, 1 μg/ml Pepstatin A, 0.5 mM PMSF, 25 mM NaF)^90^ and resolved in 8% SDS-PAGE.

### Statistical analysis

Data are represented as mean ± standard error of the mean (SEM) in XY graphs. For all other representations, whiskers and boxes were used, and the result of each biological replicate is represented in the graphs as a single dot. Circadian analysis was conducted using the R package CircaCompare^92^ or RAIN^93^ as indicated in each figure. All datasets were assessed for normality using a Shapiro-Wilk’s normality test to choose the appropriate parametric or non-parametric test. Tests used for each dataset are indicated in figure legends. Statistical differences reported throughout this work are ns p > 0.05, * p < 0.05, ** p < 0.01, *** p < 0.001 and **** p < 0.0001. The number of biological replicates is indicated in each figure or figure legend.

## Acknowledgements

We would like to thank all the members of the Chiu lab and Hamada lab for providing valuable comments in the development of the project. We also want to thank Dr. Yong Zhang for the PER-AID-GFP fly line and other flies provided, and Dr. Pamela Ronald for access to the confocal microscope. SHS is a Latin American Fellow in the Biomedical Sciences supported by the Pew Charitable Trusts. This work in the lab of JCC is supported by NIH R01 DK124068.

## Supplementary Information

**Table S1.** List of antibodies and dilutions used in this study.

| Antibody | Source | ID | Dilution | Reference |
| --- | --- | --- | --- | --- |
| Mouse anti-EYA | DSHB | EYA10H6 | WB: 1:1,000 IF: 1:200 | Boyle et al., Development, 1997;<br>Abrieux et al., PNAS, 2020 |
| Mouse anti-PDF | DSHB | PDF C7 | IF: 1:200 | Cyran et al., J Neurosc., 2005;<br>Cavanaugh et al., Cell, 2014 ; Hidalgo<br>et al., Trans Psych., 2021 |
| Mouse anti-BRP | DSHB | nc82 | IF: 1:50 | Wagh et al., Neuron, 2006 |
| Chicken anti-GFP | invitrogen | A10262 | IF: 1:1,000 |  |
| Mouse anti-HSP70 | Sigma-Aldrich | SAB4200714 | WB: 1:7,000 |  |
| Mouse anti-FLAG M2 | Sigma-Aldrich | M3165 | WB: 1:7,000 |  |
| Mouse anti-V5 | invitrogen | 46-0705 | WB: 1:5,000 |  |

DSHB: Developmental Studies Hybridoma Bank; WB: Western Blot; IF; Immunofluorescence.

**Figure S1.**
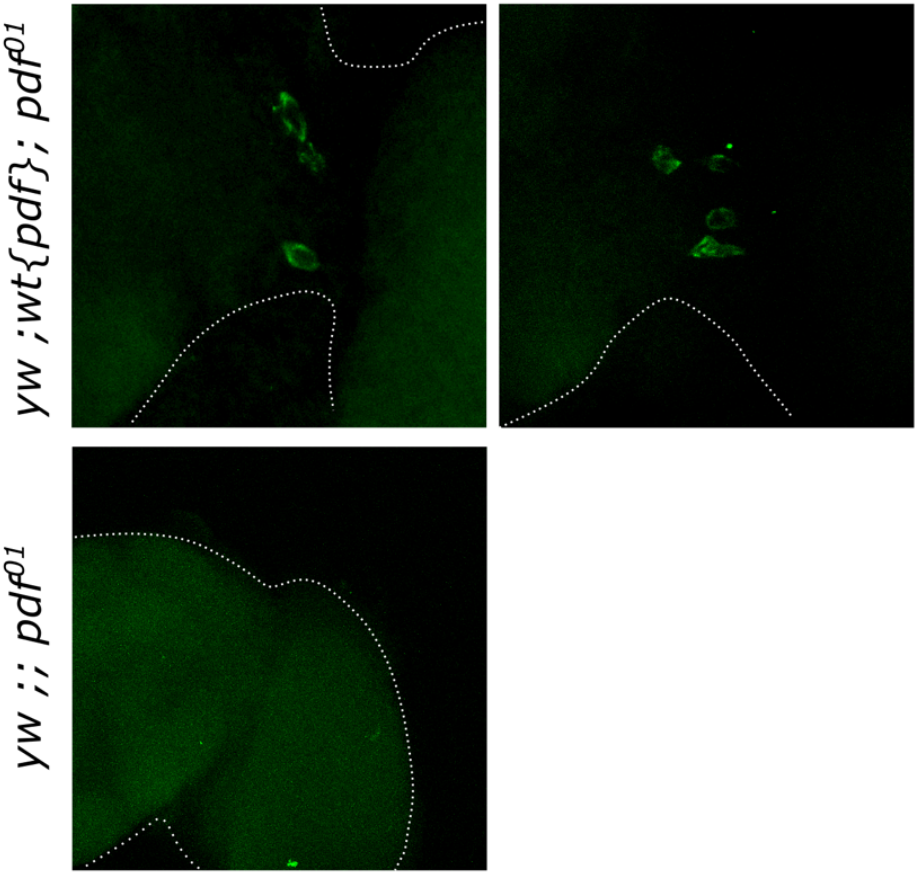
Validation of *pdf* FISH library. Examples of *pdf* mRNA detection by fluorescent *in situ* hybridization (green) in brains from *yw* ; *wt{pdf}* ; *pdf*^*01*^ genomic rescue flies (top panels) and *pdf*^*01*^ null mutants (bottom panel). As expected, no signal is observed in *pdf*^*01*^ brains.

**Figure S2.**
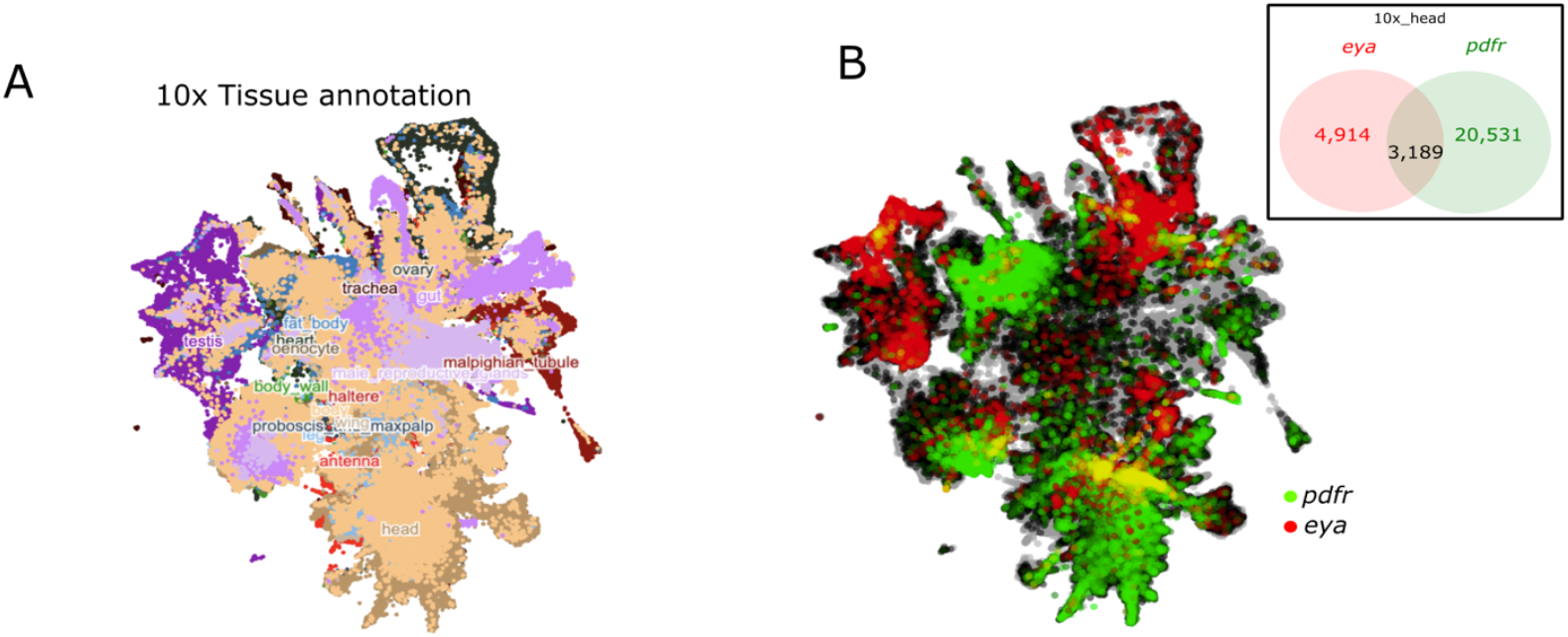
*pdfr* and *eya* expression in *Drosophila* brains. (A) All cells from the 10x relaxed dataset colored and labeled by tissue as seen by UMAP. (B) Same dataset now colored by *pdfr* (green) and *eya* (red) expression. (B, Inset) Venn diagram depicting the number of cells from the head tissue that express *eya* (red), *pdfr* (green) or both (overlap).

**Figure S3.**
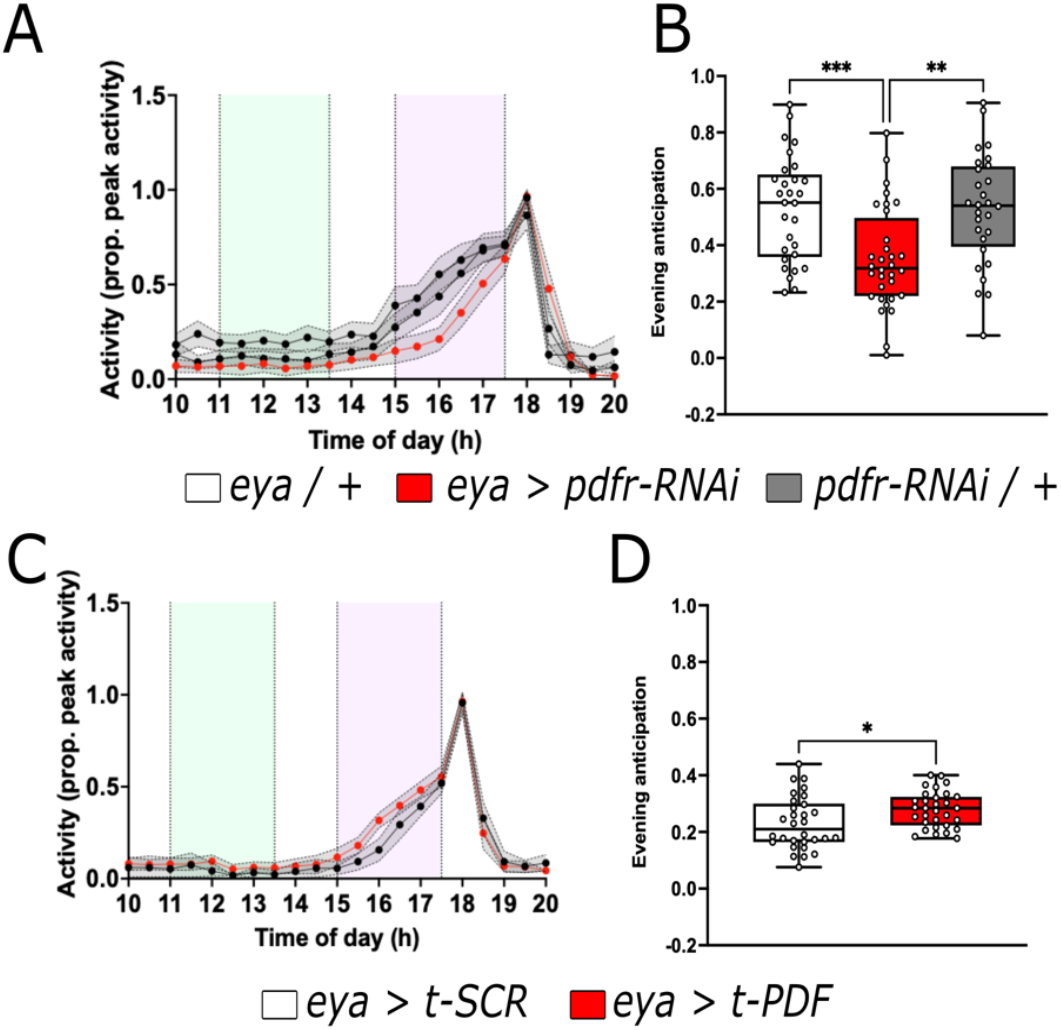
PDF modulates evening anticipation in LD through *eya+* cells. (A-B) Evening anticipation in locomotion activity of flies where the PDF receptor was knocked down in *eya*+ cells (*eya > pdfr-RNAi* red line and box) compared to controls (*eya* / + and *pdfr-RNAi* / +, black and gray lines, respectively) in 12:12 LD cycles at 25ºC. (C-D) Evening anticipation of flies where a membrane tethered version of PDF was expressed in *eya*+ cells (*eya > t-PDF* red line and box) compared to the expression of a scrambled version of the peptide (*eya > t-SCR* black line and box). Green and purple boxes in A and C denote the windows used for the calculation of the evening anticipation. Data in B were analyzed with one-way ANOVA followed by Dunnett’s multiple comparison test, unpaired t-test in D. Number of flies used were *eya / +* n = 29, *eya > pdfr-RNAi* n = 32, *pdfr-RNAi / +* n = 28, *eya > t-SCR* n = 31, *eya > t-PDF* n = 31, *eya / +* n = 30.

## Notes

### Competing Interest Statement

The authors have declared no competing interest.

